# Combined assessment of MHC binding and antigen expression improves T cell epitope predictions

**DOI:** 10.1101/2020.11.09.375204

**Authors:** Zeynep Koşaloğlu-Yalçın, Jenny Lee, Morten Nielsen, Jason Greenbaum, Stephen P Schoenberger, Aaron Miller, Young J Kim, Alessandro Sette, Bjoern Peters

**Affiliations:** Division of Vaccine Discovery, La Jolla Institute for Immunology, La Jolla, California, USA; Department of Bio and Health Informatics, Technical University of Denmark, DK Lyngby, Denmark; Instituto de Investigaciones Biotecnológicas, Universidad Nacional de San Martín, CP San Martín, Argentina; Division of Hematology and Oncology, Center for Personalized Cancer Therapy, San Diego Moore’s Cancer Center, University of California, San Diego, San Diego, California, USA; Laboratory of Cellular Immunology, La Jolla Institute for Immunology, La Jolla, California, USA; Department of Otolaryngology-Head & Neck Surgery, Vanderbilt Ingram Cancer Center, Vanderbilt University Medical Center, Nashville, Tennessee, USA; Department of Medicine, University of California, San Diego, La Jolla, California, USA

## Abstract

MHC class I antigen processing consists of multiple steps that result in the presentation of MHC bound peptides that can be recognized as T cell epitopes. Many of the pathway steps can be predicted using computational methods, but one is often neglected: mRNA expression of the epitope source proteins. In this study, we improve epitope prediction by taking into account both peptide-MHC binding affinities and expression levels of the peptide’s source protein. Specifically, we utilized biophysical principles and existing MHC binding prediction tools in concert with RNA expression to derive a function that estimates the likelihood of a peptide being presented on a given MHC class I molecule. Our combined model of Antigen eXpression based Epitope Likelihood-Function (AXEL-F) outperformed predictions based only on binding or based only on antigen expression for discriminating eluted ligands from random background peptides as well as in predicting neoantigens that are recognized by T cells. We also showed that in cases where cancer patient-specific RNA-Seq data is not available, cancer-type matched expression data from TCGA can be used to accurately estimate patient-specific gene expression. Using AXEL-F together with TGCA expression data we were able to more accurately predict neoantigens that are recognized by T cells. The method is available in the IEDB Analysis Resource and free to use for the academic community.

**Significance statement:** Epitope prediction tools have been used to call epitopes in viruses and other pathogens for almost 30 years, and more recently, to call cancer neoantigens. Several such tools have been developed, however most of them ignore the mRNA expression of the epitope source proteins. In the present study, we have, to our knowledge for the first time, developed a biophysically motivated model to combine peptide-MHC binding and abundance of the peptide’s source protein to improve epitope predictions. Our novel tool AXEL-F is freely available on the IEDB and presents a clear opportunity for predicting and selecting epitopes more efficiently.

## INTRODUCTION

Presentation of peptides on the cell surface by major histocompatibility complex (MHC) class I molecules is crucial for CD8^+^ T cell mediated immune responses, including those against viral infections and tumors. The MHC class I antigen processing and presentation pathway consists of multiple steps during which proteins are degraded into peptides, loaded on MHC class I molecules and presented on the cell surface ^1^. Recognition of these peptide-MHC complexes on the cell surface as foreign by CD8^+^ T cells prompts an immune response which can lead to the eradication of affected cells. Accurate identification of which specific peptides are presented on MHC class I has advantageous applications in diagnostics and in developing therapeutic interventions such as vaccines for infectious diseases and cancer ^2–4^.

Numerous computational tools have been developed to predict the various steps in the MHC class I antigen processing and presentation pathway (reviewed in ^5^), including prediction of proteasomal cleavage ^6, 7^, transport into the endoplasmic reticulum (ER) by transporter associated with antigen processing (TAP) ^8, 9^, peptide-MHC binding (reviewed in ^10^), and predicting the stability of the peptide-MHC complex ^11, 12^. Among these, tools predicting peptide-MHC binding have been proven to best predict immunogenic epitopes, i.e. presented peptides that are recognized by T cells ^5, 10, 13–15^. These tools generally consist of machine learning methods that have been trained with experimentally generated peptide-MHC binding data. Such experimental data is for example available in the Immune Epitope Database (IEDB) ^16^.

One drawback of peptide-MHC binding data is that only the MHC binding step of the antigen processing and presentation pathway is considered. This drawback can be overcome by using ligand elution data for training. As eluted ligands passed through the natural antigen processing and presentation pathway, the resulting elution data inherently contains valuable biological information that is not available when only peptide-MHC binding is considered ^5^. In fact, machine learning methods that have been trained on combination of peptide-MHC binding and ligand elution data outperform methods that have been trained on peptide-MHC binding data alone in predicting epitopes ^5, 10, 13, 17, 18^.

One step in the antigen processing and presentation pathway that is often ignored by epitope prediction methods is one of the earliest: mRNA expression of source proteins. Proteomic studies have previously reported correlations between protein abundance and MHC-peptide presentation ^19–21^, and more recently, it was reported that MHC-peptide presentation is strongly correlated with mRNA expression of the ligand’s source protein ^19, 22–24^, underlining the potential value of including source protein expression information into epitope predictions. In fact, Abelin et al. reported increased performance in predicting eluted ligands, and Bulik-Sullivan et al. reported increased performance in predicting immunogenic neoantigens when expression information was included in their respective machine learning models ^22, 25^.

In this study, we wanted to formally describe the interplay of peptide-MHC binding and the expression of the peptide’s source protein. We took advantage of the publicly available, highly accurate peptide-MHC binding prediction tool NetMHCpan 4.0 and developed a model that combines these predictions with the expression of the peptide’s source protein in a biophysically meaningful fashion to estimate the likelihood of the peptide being presented on a given MHC class I molecule. Our model named **A**ntigen e**X**pression based **E**pitope **L**ikelihood-**F**unction (AXEL-F) outperformed NetMHCpan 4.0 in discriminating eluted ligands from random background peptides as well as in predicting neoantigens that are recognized by T cells. AXEL-F is publicly available and free to use for the academic community at http://axelf-beta.iedb.org/axelf.

## RESULTS

### Assembly of a set of HLA class I eluted ligands and background control peptides

We wanted to assess the performance of expression level, predicted binding affinity, and their combination in distinguishing eluted ligands from a set of random background peptides. We utilized a previously published dataset of 15,090 HLA class I ligands eluted from five different HLA class I alleles ^26^. Trolle et al. isolated naturally presented ligands from HLA class I peptide complexes derived from HLA class I transfected HeLa cells (hereafter referred to as Trolle set).

Next, we generated a set of background peptides to compare the set of eluted ligands against. For each peptide in the Trolle set, 10 peptides were randomly picked from the human proteome. The lengths of the random peptides and the assignment of HLA class I alleles were chosen so that the total number of background peptides was uniformly distributed across all alleles and peptide lengths.

As expression data analysis was not included in the Trolle study, we retrieved expression data of HeLa cells from another previously published study ^27^, which used similar conditions. We used an in-house pipeline to process the raw RNA-Seq data and calculated gene expression as transcripts per million (TPM). Using the provided UniProt identifiers of the source proteins for each peptide, the corresponding TPM value was annotated to assess the expression level of the protein each peptide was retrieved from.

Next, we performed HLA class I binding predictions for each ligand and random background peptide using the NetMHCpan 4.0 algorithm ^17^. For each peptide, we retrieved predicted binding affinity provided in IC50 together with the corresponding percentile rank (BA_Rank), as well as the eluted ligand score (EL_Score) and the corresponding percentile rank (EL_Rank). The complete dataset is provided in **Supplemental Table S1**.

It is widely known ^17^ that the NetMHCpan EL_Rank and EL_Score predictions generally perform better in predicting eluted ligands than predicted IC50. The neural network that performs these EL predictions has been trained on eluted ligand data and is thus more capable of capturing eluted ligands. However, as the EL_Score is an output score of the neural network architecture, it is an abstract value and cannot be directly translated into the biological context of our model of peptide presentatio. IC50 values are defined as the concentration that inhibits 50% binding of a labeled reference peptide, and, if the assay is performed under appropriate conditions, the log(IC50) values are proportional to binding free energies. Low IC50 values correspond to peptides binding with high affinities. We therefore chose to use the predicted IC50 as a measure of peptide-HLA binding for the following analyses and provide additional analyses based on EL_Rank and BA_Rank as supplemental figures.

### HLA class I eluted ligands originate from highly expressed genes and are predicted good HLA binders

We compared expression levels of the genes from which eluted ligands originated to expression levels of the genes from which background peptides were retrieved. We analyzed each of the five alleles separately (**Figure 1A**). As expected, this analysis showed that ligands are expressed at significantly higher levels than random background peptides (p < 2.2e^−16^, Wilcoxon Test). A total of 91% of ligands were expressed above a TPM of 10, while only 48% of background peptides were expressed at this level. These results confirm that MHC I eluted ligands are preferentially derived from abundant proteins, as previously reported ^22^.

**Figure 1.**
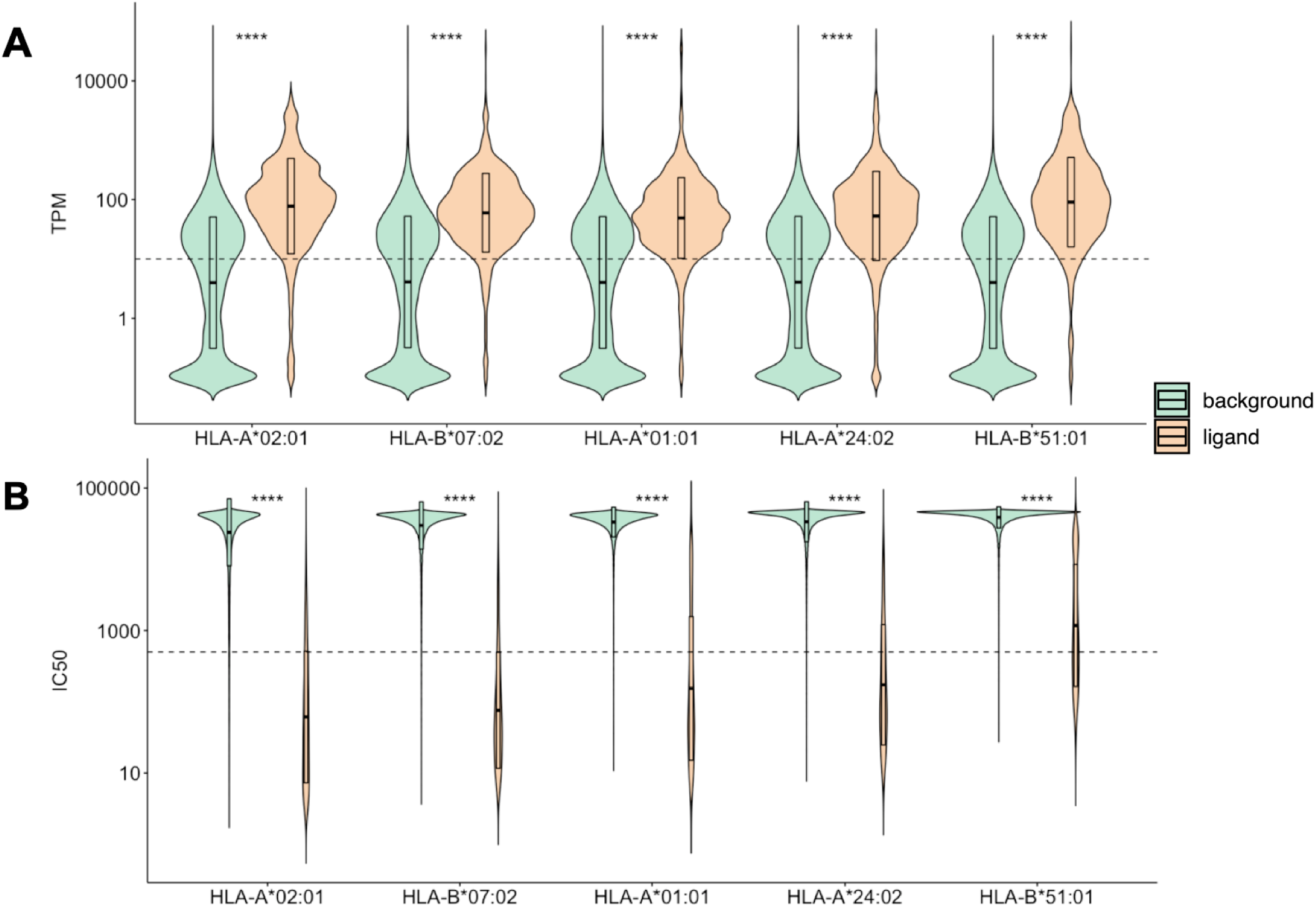
HLA class I eluted ligands originate from highly expressed genes and are predicted good HLA binders. The quartile ranges and density of TPM (A) and predicted IC50 (B) values are displayed for the five alleles included in the dataset. Ligands (displayed in tan) are expressed at significantly higher levels than random background peptides (displayed in green) and are predicted to bind at significantly higher levels (p < 2.2e^−16^, Wilcoxon Test). Dashed lines indicate TPM 10 and IC50 500 nM, respectively.

To explore how predicted HLA binding affinity of eluted ligands compare to the background peptides, we compared predicted IC50 values of eluted ligands to background peptides separately for each of the five alleles (**Figure 1B**). The majority of the background peptides (99%) were not predicted to bind (IC50 > 500 nM). In contrast, eluted ligands were predicted to bind at significantly higher levels (p < 2.2e^−16^, Wilcoxon Test) and 75% of ligands were predicted to bind their restricting HLA (IC50 < 500 nM). Similar results were obtained when this analysis was performed based on BA_Rank and EL_Rank (**Supplemental Figure S1**).

As a next step, we wanted to further investigate the relationship and interplay between HLA binding of a ligand and the expression of its source protein. We separated the binding affinity and TPM values in our dataset into ranges to create a 2-dimensional matrix with the TPM on the x-axis and the IC50 on the y-axis (**Figure 2, Supplemental Figure S2**). Each peptide was then assigned to a cell in this matrix according to its IC50 and TPM values. For each cell, we then determined the percentage of ligands among all peptides that fall into the corresponding IC50 and TPM ranges. Visual inspection of the resulting matrix revealed that certain IC50 and TPM ranges were enriched for eluted ligands, namely the part of the matrix that represents high binding affinity and substantial expression, as already discussed above. The matrix however showed additional interplay between IC50 and TPM: ligands with lower expressed source proteins seemed to bind HLA strongly while ligands that were not able to strongly bind HLA seemed to be derived from highly expressed source proteins. When we compared the IC50 values of ligands derived from the top 10% expressed source proteins to ligands derived from the bottom 10%, we found that ligands derived from low abundance proteins bind HLA significantly better than ligands derived from high abundance proteins (p-value < 2.2e-16, Wilcoxon test, **Supplemental Figure S3**).

**Figure 2.**
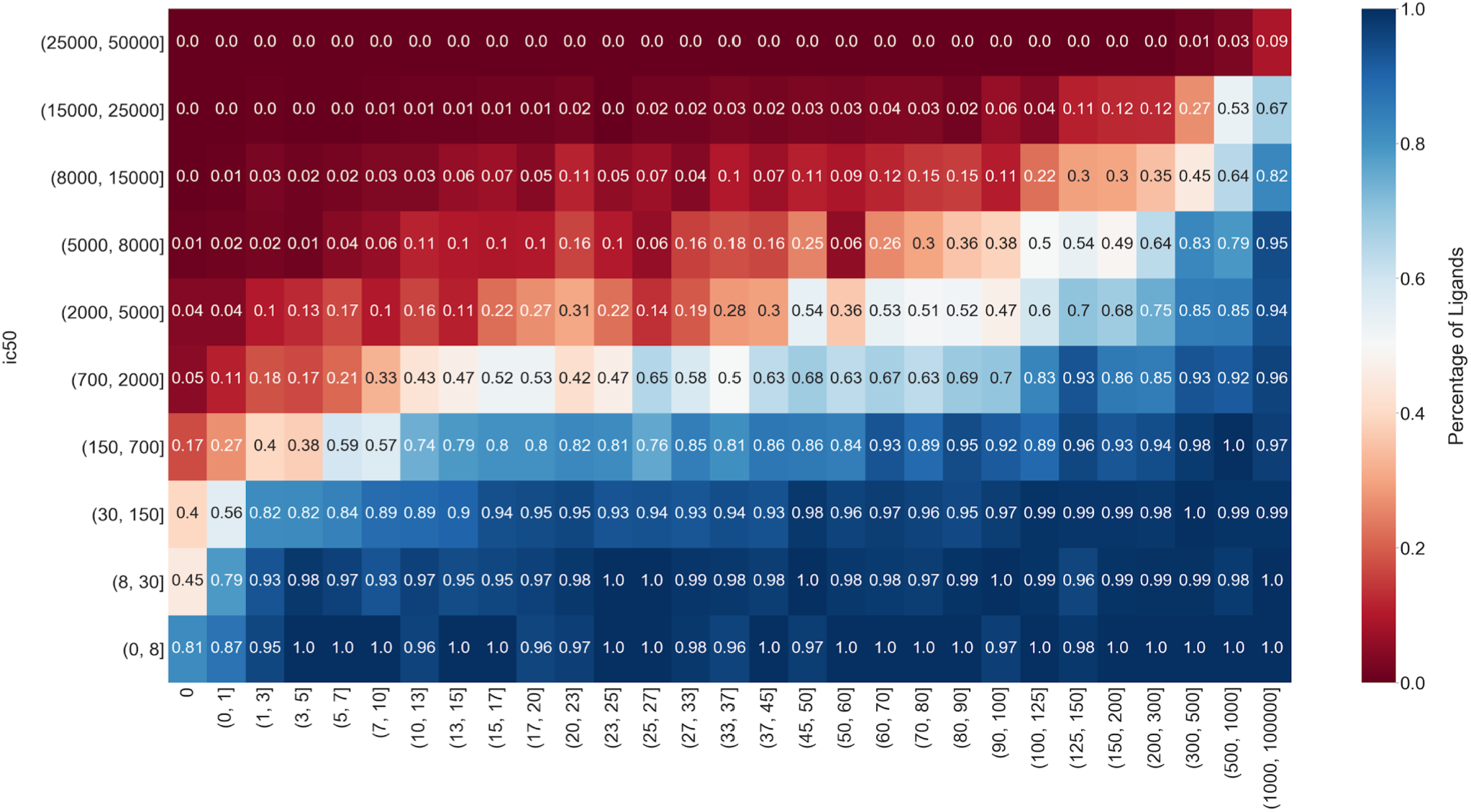
Interplay between HLA binding of eluted ligands and expression of their source Proteins. The binding affinities and TPM values were separated into ranges to create a 2-dimensional matrix with the TPM on the x-axis and the IC50 on the y-axis. Each peptide was assigned to a cell in this matrix according to its IC50 and TPM values. For each cell, the percentage of ligands among all peptides that fall into the corresponding IC50 and TPM ranges was determined and the cell was colored accordingly.

These observations, which closely mimic those of others ^22^, indicated that HLA binding of a ligand and the expression of its source protein might compensate for each other. Abundant expression of a source protein will generate more peptides which in turn might enhance the chances of these abundant peptides to bind HLA even if their HLA binding capacity is weak, simply by being available in high numbers. Conversely, a peptide with high binding affinity might still be presented on HLA even if is not abundantly expressed, by outcompeting other more abundant peptides available for HLA binding.

### HLA binding and expression level are independent predictors of HLA class I eluted ligands

Having established that eluted ligands are highly expressed and are predicted good binders, we analyzed the predictive performance of these metrics in distinguishing ligands (positives) from background peptides (negatives). We considered all four metrics provided by NetMHCpan as well as the TPM of the source protein as a measure of expression and performed a Receiver Operating Characteristics (ROC) analysis to assess prediction performance in terms of the Area Under the ROC Curve (AUC) as well as partial AUC at 10% false positive (pAUC). All NetMHCpan metrics were excellent predictors of eluted ligands: all AUC and pAUC values were above 0.99. With an AUC value of 0.812 and a pAUC of 0.629, TPM alone was also a good predictor for eluted ligands (**Figure 3)**.

**Figure 3.**
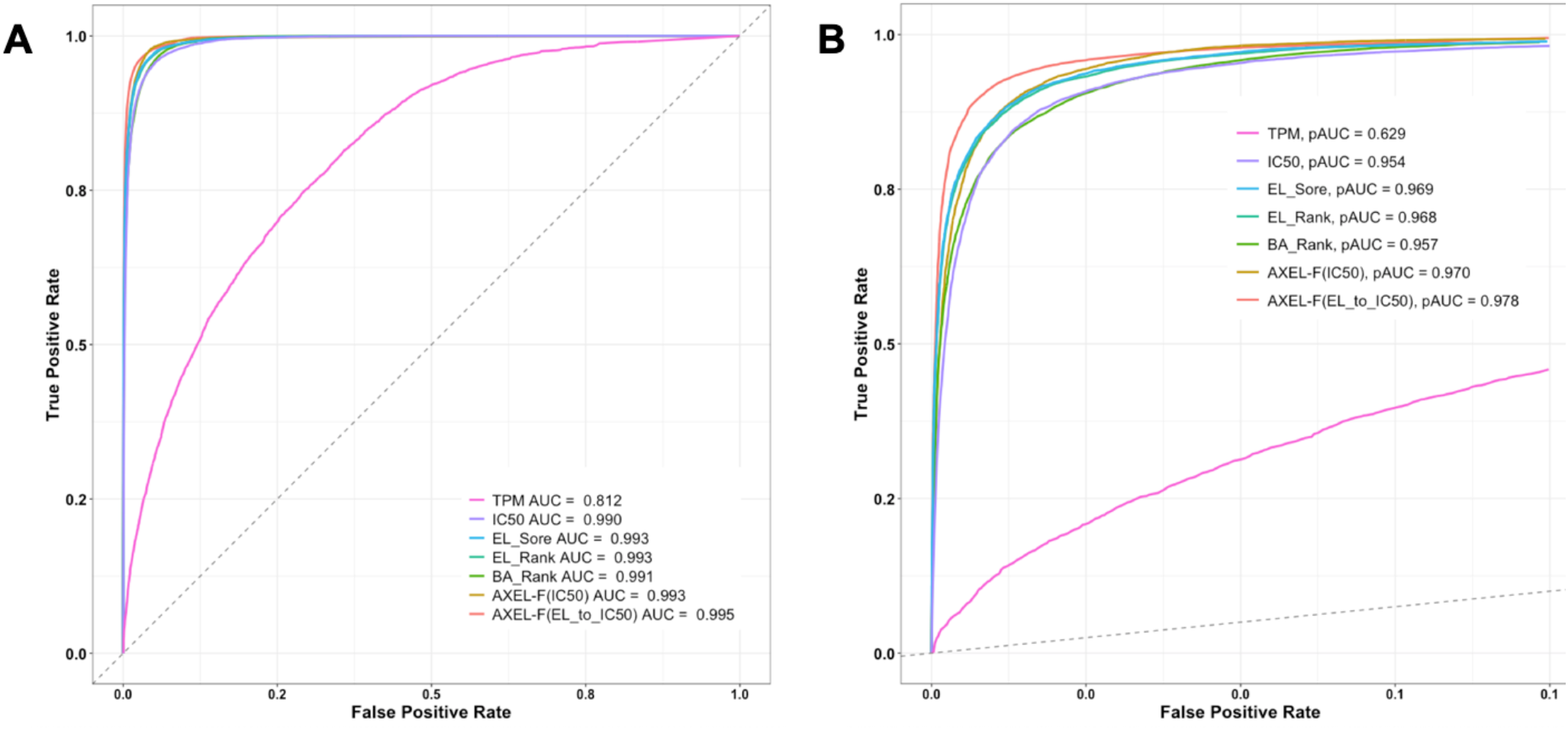
Performance of different predictors in identifying HLA class I eluted ligands. Receiver Operating Characteristics (ROC) curves (A) and ROC curves at 10% false positive rate (B) for different NetMHCpan predictors, TPM and AXEL-F scores are displayed.

### Integrating HLA binding of the ligand and expression of its source protein using a Boltzmann distribution

To integrate HLA binding capacity of the ligand and the expression of its source protein into a function to more accurately predict ligand elution, we first applied a naive approach to combine HLA binding and expression by simply assigning a poor predictive value to each peptide that was derived from a non-expressed source protein (TPM = 0). This approach was based on the biological assumption that a peptide cannot be an eluted ligand if its source protein is not expressed. We considered all peptides from the Trolle set and the background peptides and assigned each peptide the worst possible prediction score if the corresponding source protein was not expressed. A ROC analysis was performed for each metric provided by NetMHCpan and corresponding AUC values summarized in **Supplemental Table S2** clearly indicated that this naive method does not improve predictive performance across NetMHCpan predictions.

Next, we tested a more complex model to capture the effect of quantitative expression differences and combine them with peptide-HLA binding using the Boltzmann distribution which is often used to describe biophysical systems ^28^. The Boltzmann distribution is a probability distribution that predicts, in an ensemble of particles, the proportion of particles that will be in a certain state with a specific energy ^29^. This function can be adapted to describe peptide presentation, as we want to detect, among all available peptides, the ones that are in a state bound to HLA with a specific binding free energy. In this context, the number of all available peptides of a certain species was considered proportional to the expression values of its source protein (TPM) and the binding free energy can be inferred from binding affinities (IC50). Combining these considerations with the Boltzmann distribution function yields:

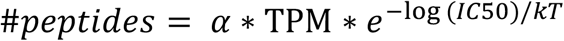

where α is a scaling factor for TPM and kT is a scale that mimics the product of the Boltzmann's constant k and the thermodynamic temperature T, as adapted from the original Boltzmann distribution function. To account for the detection limit of RNA-Seq, we additionally introduced a parameter minTPM and modified the function to select for the higher value between minTPM and the input TPM value:

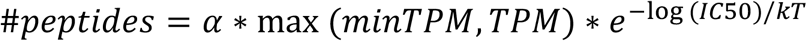

This function will estimate the number of peptides for a given species that are bound to MHC. We want to know the likelihood of finding at least one of these peptides when performing a mass spectrometry experiment and/or when a T cell scans a cell:

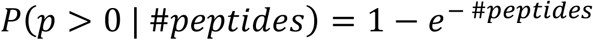

Our final model estimates, for a given peptide with IC50 and TPM value, its likelihood of being presented on HLA and being an epitope. We named this model AXEL-F, standing for **A**ntigen e**X**pression based **E**pitope **L**ikelihood-**F**unction.

Next, we used the Trolle dataset to identify the optimal value for the three free parameters of AXEL-F (α, kT, and minTPM), based on the predictive performance measured by AUC. We performed 10 iterations of 5-fold cross-validation to avoid overfitting the parameters and also trained the parameters separately for the 5 alleles as well as for the complete dataset. For each parameter, we used the median value over all runs of cross-validation. The parameters obtained in this way fell into a consistent range between the 5 subsets corresponding to the 5 alleles in the Trolle set (**Table 1**).

**Table 1.** Fitted parameters and obtained AUC values.

| Allele | $\alpha$ | kT | minTPM | AUC AXEL-F | AUC IC50 | AUC EL_rank |
| --- | --- | --- | --- | --- | --- | --- |
| HLA-B*07:02 | 1.267 | 0.133 | 0.567 | 0.996 | 0.994 | 0.995 |
| HLA-A*01:01 | 1.296 | 0.119 | 0.533 | 0.995 | 0.993 | 0.994 |
| HLA-A*02:01 | 1.173 | 0.215 | 0.612 | 0.992 | 0.987 | 0.993 |
| HLA-A*24:02 | 1.267 | 0.133 | 0.567 | 0.996 | 0.993 | 0.995 |
| HLA-B*51:01 | 1.292 | 0.123 | 0.521 | 0.991 | 0.987 | 0.982 |
| **Complete dataset** | **1.296** | **0.131** | **0.523** | **0.993** | **0.990** | **0.993** |

AXEL-F outperformed IC50 consistently for all 5 alleles as well as for the complete dataset (p-value = 0.002, DeLong's test) as shown by increased AUC values (**Table 1**). AUC values for AXEL-F scores were also slightly higher than those for EL_Rank for all alleles except HLA-A*02:01, the increase in AUC however was only significant for HLA-B*07:02 (p < 0.02, DeLong's test). AXEL-F and EL_Rank were on par when applied on the complete dataset. As the gain in performance was significantly higher when parameters were trained on the complete dataset (p-value < 2.2e-16, DeLong’s test), we decided to present subsequent analyses for the complete dataset only.

As mentioned and shown above, EL_Rank predictions generally perform better in predicting eluted ligands than predicted IC50. However, as the EL_Rank is an abstract output value of the neural network architecture it and cannot be directly translated into the biological context of our model. We still wanted to take advantage of the superior predictive performance of El_Rank and incorporate it into our model. In order to achieve this, we mapped each EL_Rank to a corresponding IC50 value by comparing the percentile ranks of the two metrics. We named this new metric EL_to_IC50 and used it as an input for AXEL-F instead of the IC50 values we used earlier, and again used the Trolle dataset to train the parameters α, kT and minTPM. The values we obtained were very similar to those obtained when using IC50: α was fit to 1.233, kT to 0.156, and minTPM to 0.567. We were able to additionally further boost the performance of AXEL-F and achieved an AUC of 0.995 and a pAUC of 0.978 (**Table 2, Figure 3**). AXEL-F (EL_to_IC50) significantly outperformed AXEL-F (IC50) as well as all NetMHCpan predictions (p-value < 2.2e-16, DeLong’s test).

**Table 2.** Overview of AUC and pAUC values for Trolle and Abelin datasets.

|  | **Trolle** |  | **Abelin** |  |
| --- | --- | --- | --- | --- |
|  | AUC | pAUC | AUC | pAUC |
| **TPM** | 0.812 | 0.629 | 0.761 | 0.605 |
| **IC50** | 0.990 | 0.954 | 0.965 | 0.930 |
| **BA_Rank** | 0.991 | 0.957 | 0.968 | 0.946 |
| **EL_Score** | 0.993 | 0.969 | 0.974 | 0.950 |
| **EL_Rank** | 0.993 | 0.968 | 0.973 | 0.953 |
| **AXEL_F (IC50)** | 0.993 | 0.970 | 0.973 | 0.940 |
| **AXEL-F (EL_to_IC50)** | **0.995** | **0.978** | **0.978** | **0.955** |

### Model evaluation on an independent dataset of eluted ligands

We next validated our results with an independent dataset ^22^ of eluted ligands from mono-allelic B cells transduced with a single HLA class I molecule. The dataset contained 26,089 eluted ligands from 16 HLAs, with only four overlapping with those present in the Trolle dataset. Abelin et al also performed RNA-Seq and provided the data in the form of TPM values (GSE93315). We again generated a background dataset and performed NetMHCpan predictions for all 260,890 peptides. The complete dataset is provided in **Supplemental Table S3**.

We calculated AXEL-F likelihood scores by using both, IC50 and EL_to_IC50, as inputs and compared the performance to NetMHCpan predictions in discriminating eluted ligands from the random background set. The AUC and pAUC results summarized in **Table 2** and the ROC curves shown in **Supplemental Figure S4** indicate that AXEL-F likelihood scores significantly outperformed IC50 and BA_Rank when likelihood scores are calculated using IC50 (AXEL-F (IC50), p-value < 2.2e-16, DeLong’s test). When likelihood scores were calculated using EL_to_IC50 (AXEL-F (EL_to_IC50)) it also significantly outperformed EL_Score and EL_Rank (p-value < 2.2e-16, DeLong’s test).

Overall, these data showed that AXEL-F outperformed all NetMHCpan predictions alone on both the original Trolle dataset, and a comprehensive independent ligand elution dataset. These results highlight the robustness of our model and the biological relevance of integrating HLA binding and expression of the source protein of a given peptide in predicting whether the peptide is an eluted ligand or not.

### Model evaluation on a dataset of experimentally validated cancer neoantigens

We next wanted to analyze how our model AXEL-F performed in predicting epitopes, specifically cancer neoepitopes that arise from somatic mutations. We utilized a previously published study by Parkhurst et al ^30^ that reported immunogenicity screening results of neoantigens from 75 patients with various gastrointestinal cancers. The group performed whole-exome sequencing to detect somatic mutations and determined which neoantigens were recognized by tumor-infiltrating-lymphocyte cultures ^30^. The results were provided as a supplemental table to the study, listing all tested neoantigens and corresponding screening results (CD4^+^ and/or CD8^+^ or negative, (Supplementary 3 in ^30^).

As our study is based on HLA class I predictions, we only considered the 54 neoantigens that were recognized by CD8^+^ T cells and the 7,529 peptides that were not recognized at all. We further filtered the dataset by only retaining peptides for which RNA-Seq information was provided, which resulted in a final set of 46 patients with 28 recognized neoantigens and 1,298 peptides that were not recognized. We named this dataset the NCI dataset and wanted to compare the performance of AXEL-F to NetMHCpan predictions alone in distinguishing immunogenic neoantigens (positives) from peptides that were not recognized by tumor infiltrating lymphocytes (negatives).

To do so, we first performed NetMHCpan predictions on the complete dataset and found that both, IC50 and EL_Rank could discriminate immunogenic neoantigens with AUC values of 0.698 and 0.688, respectively (**Table 3)**. The RNA-Seq information that was provided with the NCI dataset included read depth at the position of the mutation (tumor_rna_depth), the number of reads supporting the mutation (tumor_rna_alt_reads), and the variant allele frequency (tumor_rna_alt_freq). Unfortunately, TPM values were not provided as part of the dataset. As a first analysis, we assessed the predictive performance of these RNA metrics in predicting immunogenic neoantigens and found that all three metrics had some predictive value (**Table 3 and Supplemental Figure S5**). With an AUC of 0.642, the number of reads supporting the mutation (tumor_rna_alt_reads) had the best performance among the three RNA metrics. We presumed that the tumor_rna_alt_reads can be used as a proxy for describing the expression of mutated transcripts and used this metric instead of a TPM together with the EL_to_IC50 to calculate AXEL-F likelihood scores. In fact, the AUC values for AXEL-F scores that were obtained this way (AXEL-F (tumor_rna_alt_reads)) were higher than those of both NetMHCpan predictions as well as the tumor_rna_alt_reads alone, with an AUC of 0.722 (**Table 3**). The difference in AUC however, was not significant (DeLong’s test).

**Table 3.**
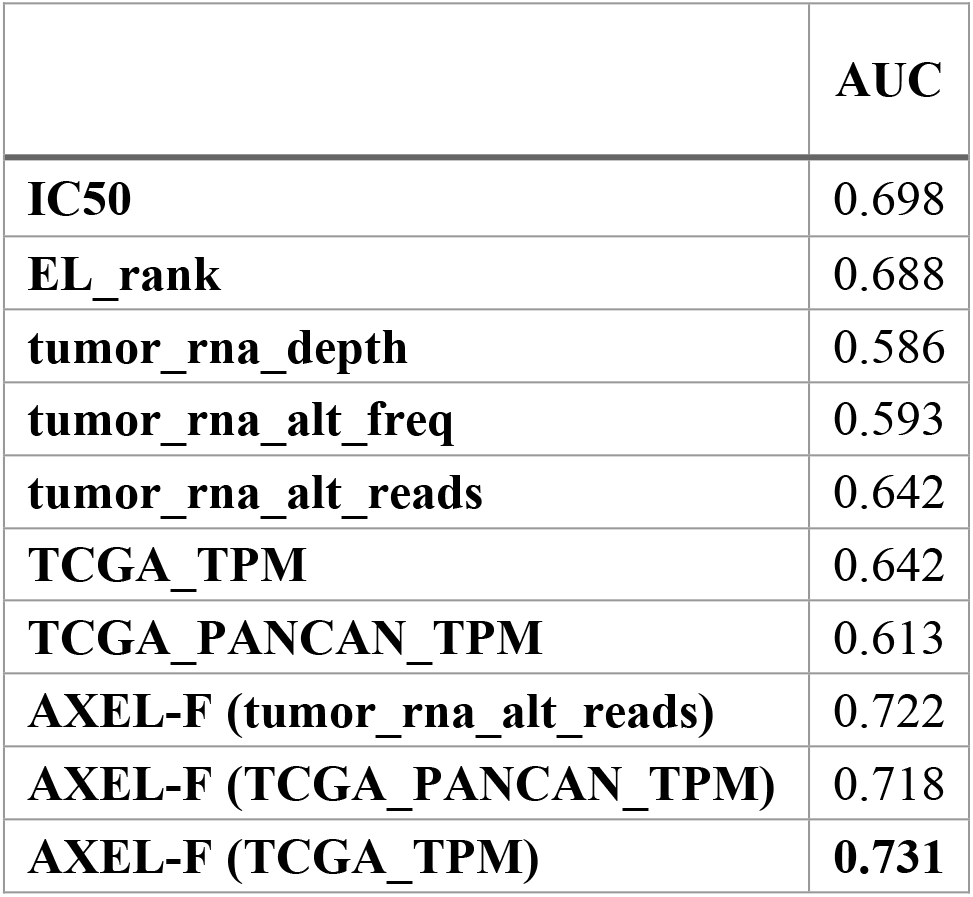
Overview of AUC values in predicting immunogenic neoantigens from the NCI dataset.

|  | AUC |
| --- | --- |
| **IC50** | 0.698 |
| **EL_rank** | 0.688 |
| **tumor_rna_depth** | 0.586 |
| **tumor_rna_alt_freq** | 0.593 |
| **tumor_rna_alt_reads** | 0.642 |
| **TCGA_TPM** | 0.642 |
| **TCGA_PANCAN_TPM** | 0.613 |
| **AXEL-F (tumor_rna_alt_reads)** | 0.722 |
| **AXEL-F (TCGA_PANCAN_TPM)** | 0.718 |
| **AXEL-F (TCGA_TPM)** | **0.731** |

### TPM values from TCGA can be used to accurately estimate gene expression in a given patient sample

As TPM values were not provided as part of the NCI dataset we wanted to analyze whether it is possible to estimate gene expression levels in a given patient sample by using expression data retrieved from The Cancer Genome Atlas (TCGA). We downloaded pre-calculated TCGA TPM values, and for each cancer type included, we calculated the median expression for each gene across all samples in the corresponding cancer type specific subset. We utilized in-house RNA-Seq data of 25 patients with 9 different cancer types and analyzed how well gene expression of these patients can be estimated by using the gene expression data retrieved from TCGA. For each patient from our in-house cohort, we matched the TPM from in-house RNA-Seq with TPM values from TCGA. This was done for each cancer type separately so that our in-house data was matched with all available cancer types from TCGA. We then analyzed, how well TCGA median TPM values correlate with the in-house RNA-Seq TPM values. This analysis showed that cancer-type matched TPM values correlate very well (Pearson’s r^2^ > 0.6, **Figure S6**).

Having established that TPM values from TCGA can be used to estimate gene expression in a given patient sample, we proceeded to utilize this approach to estimate TPM values for the NCI dataset. We matched the cancer type and gene name for each peptide in the NCI dataset and assigned the corresponding cancer type specific median TPM from TCGA. With an AUC of 0.642 this TCGA_TPM alone was as predictive for immunogenic neoantigens as tumor_rna_alt_reads alone. When we used the TCGA_TPM together with the EL_to_IC50 as inputs for AXEL-F, we achieved the best performance reaching AUC values of 0.731 (**Table 3 and Supplemental Figure S5**). Of note, when we did not match the cancer types and used TPM values calculate from the entire TCGA dataset (TCGA PANCAN) the AUC was 0.718 and thereby lower compared to when cancer types were matched.

These results underline the general applicability of our model and furthermore its potential to predict cancer neoantigens more accurately. Even when patient-specific expression data is not available, which does occur often due to the many technical challenges of RNA-Seq, it is possible to estimate the expression of neoantigen source proteins from TCGA and perform accurate predictions using AXEL-F.

## DISCUSSION

In this study, we used a biophysical model to formally describe the roles of antigen expression and peptide-MHC binding affinity in the antigen processing and presentation machinery. We hypothesized that the likelihood of a peptide being presented on HLA class I and subsequently being recognized by CD8+ T cells is dependent on both, the abundance of its source protein and its HLA binding capacity. AXEL-F, the model we developed, outperformed NetMHCpan 4.0 in discriminating eluted ligands from random background peptides as well as in predicting neoantigens that are recognized by T cells.

It had already been reported that including expression data of source proteins when training machine learning models for predicting ligand elution or neoantigen prediction can improve performance ^22, 25^. These tools however, are not publicly available. AXEL-F in contrast, is available on the IEDB Analysis Resource (IEDB-AR) and is free to use for the academic community ^31^. The IEDB-AR provides numerous computational tools focused on the prediction and analysis of B and T cell epitopes and AXEL-F is a valuable addition to these tools.

We showed that the expression level of source proteins alone is already a good predictor of ligand elution. The predictive value is even more pronounced in the case of neoantigens: even though patient-specific expression data was not available and we used publicly available cancer type matched expression data from TCGA, the predictive performance of this estimated TPM was almost on par with NetMHCpan predictions. Importantly, neoantigens included in the NCI dataset were not pre-filtered based on MHC binding or expression ^30^, which makes it possible to accurately compare the performance of these metrics.

Currently, many epitope prioritization algorithms only use expression data to filter out candidate neoantigens that do not meet a specified gene expression threshold ^32^. Our results indicate that the value of antigen expression levels has potentially been underestimated, and rather than using expression as a filtering step, it might be used in combination with HLA binding predictions to more efficiently identify neoantigens.

As TPM values estimate the expression of a gene, we first presumed that in the case of neoantigens, tumor_rna_alt_reads would be a better predictor than TPM as this metric specifically describes the expression of mutated transcripts. However, the predictive performance of these metrics alone was the same and when we used these metrics in our model, the model using TCGA_TPM clearly outperformed the model using tumor_rna_alt_reads. One reason for this might be that we trained our model using TPM values of source proteins of eluted ligands and the parameters would have been fitted differently when trained with a neoantigen specific dataset using tumor_rna_alt_reads. However, neoantigen datasets that provide expression details are still limited and we are unfortunately not able to further explore this approach at this point. Another complication with using tumor_rna_alt_reads to estimate the expression of mutated transcripts is that this metric directly counts the RNA-seq reads that support the mutation and is, in contrast to TPM, not normalized considering the total number of mapped reads. As more datasets become available, we will further explore how to best specifically estimate the abundance of mutated transcripts.

Due to time or financial limitations and given the many challenges of obtaining, preserving and sequencing RNA samples, RNA-Seq data is often not available in a clinical setting. Here, we have shown that expression data publicly available from TCGA can be used to effectively estimate patient-specific gene expression values. We have included TCGA expression values for 35 different cancer types in the current implementation of AXEL-F and plan to extend the available expression datasets to cell lines and different cell types.

Many factors during an RNA-Seq experiment can have an effect on gene expression and thus on the TPM of a specific gene. In our model we introduced the parameter minTPM to accurately capture lowly expressed transcripts. This was necessary because RNA-Seq has a detection limit and additional bias is imposed by only working with a single RNA-Seq sample, as it is mostly not feasible to generate biological and technical replicates in a clinical setting. The optimal value we obtained for minTPM was 0.567, indicating that every transcript that generated a ligand should have been expressed at least with a TPM of 0.567 at some point in the cell cycle. Biologically speaking, there is no definitive minimum TPM value above which a gene can be considered expressed. Technically speaking however, a TPM cutoff is often utilized to select genes that are considered expressed. The EMBL-EBI Expression Atlas for example, uses a default minimum expression level of 0.5 TPM ^33^. This value is very close to the minTPM value that our training determined and thus supports the biological relevance of this value.

How the interplay between source antigen expression and peptide-MCH binding can be utilized to more efficiently predict viral epitopes remains an open question. During infection, viruses hijack host cells to express genes necessary for virus propagation. Which genes are expressed and at what level depends on several factors ^34^. The kinetics of the viral infection play an important role as different genes are expressed during different stages of the viral infection ^35^, and importantly, many viruses subvert the MHC processing and presentation pathway at later stages of the infection. Hence, to include source antigen expression data for viral epitope prediction it would be necessary to know the kinetic class of the antigen of interest. The genes in each kinetic class, however, are different for each viral family and are not well known for many viruses. Additionally, viral gene expression varies significantly among genetically identical cells and the source of these variations is still not well understood ^34, 36^. Given these complications in capturing viral gene expression, AXEL-F is currently not suitable to predict viral epitopes.

Finally, we did not address prediction of MHC class II restricted epitopes presented to CD4^+^ T cell, which play an important role in autoimmunity and antitumor immunity. While MHC class I binding peptides are mainly derived from endogenous proteins, peptides binding MHC class II are mainly derived from extracellular proteins. Our model needs to be adjusted to describe the MHC class II antigen presentation pathway, and the cellular location of the source antigen might be one variable to consider. Unfortunately, the quality of eluted ligands from MHC class II is still lacking as ligand elution experiments from MHC class II are more complex when compared to MHC class I. Due to the open binding groove of MHC class II, ligands are variable in length and it is difficult to deconvolute multi-allelic ligand data. Recently, more computational methods to accurately deconvolute multi-allelic ligand data are becoming available ^37 38, 39^ and also more ligands eluted from mono-allelic cell lines are being published. This will help to retrieve quality datasets that we can use to train a model for MHC class II presentation.

Taken together, we have, to our knowledge, for the first time developed a biophysically motivated model to combine peptide-MHC binding and abundance of the peptide’s source protein to improve epitope predictions. AXEL-F is freely available and should be useful for predicting and selecting epitopes more efficiently.

## MATERIALS AND METHODS

### Training Dataset

For our initial analysis and as the training set for model development, we used a previously published dataset of 15,090 HLA class I ligands eluted from five different HLA class I alleles: HLA-A*01:01, HLA-A*02:01, HLA-A*24:02, HLA-B*07:02, and HLA-B*51:01 ^26^. We downloaded this dataset from the IEDB under the accession number 1000685 (http://www.iedb.org/subID/1000685). The length of the ligands in this set ranged from 8 to 14 residues. Eluted ligands were retrieved from 4,831 different source proteins for which UniProt identifiers were also provided.

As expression data was not included in the Trolle study, we retrieved expression data of HeLa cells from another previously published study ^27^. We downloaded raw read data from the Gene Expression Omnibus database under accession number GSM3899456 and used an in-house pipeline to process the raw RNA-Seq data and calculate gene expression as transcripts per million (TPM).

### Validation dataset of eluted ligands

We retrieved a second dataset of eluted ligands to validate our findings ^22^. The dataset was provided as Supplementary tables and contained 26,089 eluted ligands from 16 HLA class alleles: HLA-A*01:01, HLA-A*02:01, HLA-A*02:03, HLA-A*02:04, HLA-A*02:07, HLA-A*03:01, HLA-A*24:02, HLA-A*29:02, HLA-A*31:01, HLA-A*68:02, HLA-B*35:01, HLA-B*44:02, HLA-B*44:03, HLA-B*51:01, HLA-B*54:01, and HLA-B*57:01. Abelin et al also performed RNA-Seq and provided the data in the form of TPM values from four cell lines (GSE93315). We averaged TPM values from those four cell lines.

### Background data generation

To compare the set of eluted ligands against, we generated sets of random background peptides. For each peptide in the training or validation dataset, 10 peptides were randomly picked from the human proteome. The lengths of the random peptides and the assignment of HLA class I alleles were chosen in a way that the total number of background peptides was uniformly distributed across all alleles and peptide lengths.

### Dataset of validated immunogenic neoantigens

We used data from a previously published study by Parkhurst et al ^30^ that reported immunogenicity screening results of neoantigens from 75 patients with various gastrointestinal cancers. The group performed whole-exome sequencing to detect somatic mutations, transfected autologous dendritic cells with tandem minigenes encoding these mutations, and determined which neoantigens were recognized by tumor-infiltrating-lymphocyte cultures ^30^. The results were provided as a supplemental table to the study, listing all tested neoantigens and corresponding screening results (CD4+ and/or CD8+ or negative, (Supplementary 3 in ^30^).

The peptides provided in this dataset were mainly 29mer peptides with the mutated residue located in the center of the peptide. When performing NetMHCpan predictions, we considered all contained 8-12mer peptides and all HLA class I alleles provided for the corresponding patient. For each peptide we then assigned the best IC50 and the best EL_rank predictions among all its k-mer and HLA combinations.

### TCGA Analysis

We downloaded pre-calculated TPM values for the TCGA Pan-cancer cohort from UCSC Xena datapages ^40^. For each of the 35 cancer types included, we calculated the median expression for each gene across all samples. We utilized in-house RNA-Seq data of 25 patients with 9 different cancer to analyze how well patient-specific gene expression can be estimated by using the gene expression data obtained from TCGA. For each patient from our in-house cohort, we matched the TPM from in-house RNA-Seq with the calculated cancer type specific median TPM values from TCGA. We then analyzed, how well TCGA median TPM values correlated with patient-specific TPM values.

### HLA class I Binding Predictions

NetMHCpan version 4.0 as hosted on the IEDB Analysis Resource (IEDB-AR) was used to perform binding predictions ^17, 31^.

### Transforming EL_Rank to IC50 values

EL_Score and EL_Rank are output values of the neural networks the NetMHCpan method consists of. These values are abstract and cannot be directly used in the biological context of our model like IC50. We therefore translated the EL_Rank values to IC50 values by comparing the percentile ranks of the two metrics in the Trolle dataset. To do so, we first calculated the global percentile rank of each IC50 value within the Trolle set. We then defined an interpolation function that maps each of these percentile ranks to the corresponding IC50 value. This interpolation function was then used to map each EL_Rank to IC50 values to obtain our new metric EL_to_IC50. The same interpolation function based in the Trolle dataset was used to calculate AXEL-F scores for the validation datasets Abelin and NCI and the function is implemented as part of the AXEL-F method.

### Performance evaluation

The R package pROC was used for performing ROC analysis and calculating AUC and pAUC values, packages plotROC and ggplot 2 were used to plot ROC curves.

### Model Training

We used the R function optim that implements an optimization method based on Nelder–Mead. The paramters α, minTPM and kT were fitted to maximize the AUC value for predicting eluted ligands in the Trolle dataset. To avoid overfitting, we performed 5-fold-cross-validation: the dataset was split randomly into 5 parts using R package caret, the optimization was performed on 4/5 of the data and tested on the remaining 1/5 of the data. This was done 10 times and the median of the fitted parameters were used for α, minTPM and kT.

